# Inference of Brain States under Anesthesia with Meta Learning Based Deep Learning Models

**DOI:** 10.1101/2021.12.05.471326

**Authors:** Qihang Wang, Feng Liu, Guihong Wan, Ying Chen

## Abstract

Monitoring the depth of unconsciousness during anesthesia is useful in both clinical settings and neuroscience investigations to understand brain mechanisms. Electroencephalogram (EEG) has been used as an objective means of characterizing brain altered arousal and/or cognition states induced by anesthetics in real-time. Different general anesthetics affect cerebral electrical activities in different ways. However, the performance of conventional machine learning models on EEG data is unsatisfactory due to the low Signal to Noise Ratio (SNR) in the EEG signals, especially in the office-based anesthesia EEG setting. Deep learning models have been used widely in the field of Brain Computer Interface (BCI) to perform classification and pattern recognition tasks due to their capability of good generalization and handling noises. Compared to other BCI applications, where deep learning has demonstrated encouraging results, the deep learning approach for classifying different brain consciousness states under anesthesia has been much less investigated. In this paper, we propose a new framework based on meta-learning using deep neural networks, named Anes-MetaNet, to classify brain states under anesthetics. The Anes-MetaNet is composed of Convolutional Neural Networks (CNN) to extract power spectrum features, and a time consequence model based on Long Short-Term Memory (LSTM) Networks to capture the temporal dependencies, and a meta-learning framework to handle large cross-subject variability. We used a multi-stage training paradigm to improve the performance, which is justified by visualizing the high-level feature mapping. Experiments on the office-based anesthesia EEG dataset demonstrate the effectiveness of our proposed Anes-MetaNet by comparison of existing methods.

## I. Introduction

Providing the right dose of anesthetic drugs to a patient is of paramount importance for safely and humanely performing most surgical and many nonsurgical procedures [1]. Underdoses of anesthetic drugs will make patients wake up during surgery, while an overdose of drugs increases the risk of post-operative delirium [2] [3]. The inference of brain states provides a unique opportunity for precise administration of drugs on different subjects based on real-time closed-loop anesthesia delivery system [4]. According to the Richmond Agitation-Sedation Scale (RASS), anesthesiologists mostly estimate the depth of anesthesia (DoA) manually at intervals; however, this would require considerable attention with a complicated stimuli-response procedure during the surgery [5]. Measurements through the observation of heart rate, breathing pattern, blood pressure and other factors, have been used to measure DoA [6]; however, these physiological signals are the secondary measurements of DoA, showing large cross-subject variability that requires significant experience from anesthesiologists to decode the DoA. In recent years, EEG as an invaluable modality of recording brain activity has attracted more and more attention among anesthesiologists [7]. Compared to the secondary measurements, EEG signal is a direct measurement for brain states as the brain cognitive process relies on communications between neuronal populations through electrical signal [8]. Importantly, it has been found that DoA is associated with EEG signatures given different drugs [3] [9].

### Existing work

Numerous EEG-based monitoring frameworks for the DoA have been developed using statistical or machine learning models [10]–[12]. Specifically, Shalbaf *et al.* monitored the DoA by extracting entropy features from EEG and using artificial neural networks for classification of DoA [13]. Peker *et al.* estimated the DoA by combining ReliefF feature selection and random forest algorithm [14]. Jahanseir *et al.* estimated the DoA with a multi-output least-square support vector regression method [15]; Saadeh *et al.* assessed the DoA with a machine learning fine decision tree classifier for DoA classification of 4 states (deep, moderate, and light DoA versus awake state) [16]. In addition to the above classical machine learning models, recently, many researchers employed deep learning as classification models for monitoring the DoA. Compared to the traditional machine learning models, deep learning has demonstrated outstanding performance on many computer vision and natural language processing tasks among other interesting applications [17]–[21]. As for the DoA task, only a few researches based on deep learning have been conducted. For instances, Park *et al.* proposed a real-time DoA monitoring system based on a convolutional neural network framework [22]; Li *et al.* introduced a novel method based on hybrid features including sample entropy, permutation entropy, spectra, and alpha-ratio and performed classification using recurrent neural network [23]; Afshar *et al.* developed a combinatorial deep learning structure involving convolutional neural networks, bidirectional long short-term memory, and an attention layer to estimate the DoA precisely from EEG signals [24].

### Knowledge gap and our contribution

Although previously proposed deep-learning methods outperform the existing classical machine learning algorithms, these studies are designed to explore the anesthetic data collected from the hospital-based environment. Compared to hospital-based anesthesia (HBA), office-based anesthesia (OBA) is conducted in an outpatient setting. The advantages of OBA include easy scheduling, cost containment, and improved patient privacy, etc. However, it also has some disadvantages, namely slow recognition or response to emergent situations and lack of experience intensively monitoring of drug administration, which would result in a high level of motion artifacts and low signal to noise ratio (SNR) in the collected EEG data. In other words, the data from OBA are more challenging to analyze. However, to the best of our knowledge, there have been no previous studies that build classification models for OBA with deep learning frameworks. To bridge the research gap, in this study, we focus on developing an automated deep-learning classification framework for monitoring brain states under anesthetics in the OBA context. We use the open OBA dataset presented in Ref. [25] for explorations. However, two issues should be taken into account. The first issue we try to address is the cross-subject variability within the EEG data. The EEG signals among subjects can have high intra-class variability. Applying some signal processing methods, such as wavelet transformation [26], [27], Fourier transformation [28], [29], multitaper analysis [30] and artifacts removal algorithms [31], can reduce the impact of noise. However, there is no principled way to address the inter-subject variability. As demonstrated in the Ref. [32]–[34], convolutional neural networks (CNN) can extract features of EEG data well and have achieved outstanding performance in the hospital-based DoA classification. However, the traditional CNN are difficult to achieve a good cross-subject classification result. In other words, the direct application of CNN on the EEG data from anesthetic patients might result in unsatisfactory performance on the testing data, although the training performance can be good. Therefore, how to address the inter-subject variability under OBA becomes the main focus of this study. Secondly, given the different contexts of OBA and HBA, classifying the DoA based on OBA can be challenging due to the high level of noise in OBA. Moreover, this OBA dataset was collected under a mixed infusion of propofol, ketamine, dexmedetomidine and lidocaine, which contributes a lot to the intra-class variability. We propose to utilize the temporal dependency of spectrum features with long short-term memory (LSTM) networks to mitigate the high level of noise. Our assumption is the brain only maintains one state given a short period of time. As for subject variability, inspired by the great success of metalearning in dealing with cross-subject variability [35], we propose to incorporate the deep learning framework into the meta-learning paradigm. The essence of using meta-learning is to increase the generalization ability of the learner in multiple tasks [36]. Hence, addressing these two issues motivates us to develop a robust DoA estimation framework based on a metalearning with deep learning models. Given the unbalanced samples from different classes, we introduce a sequential classification method by first classifying the classes with better separability and then classifying the remaining classes. The experiments are conducted on the subjects with quality EEG data, and six methods are used as benchmarks to demonstrate the advantages of the proposed model.

We name the proposed network employing CNN, LSTM with sequential classification framework under the framework of meta learning as Anes-MetaNet. To summarize, our contributions can be listed as:

- We propose to use deep learning to build the classification framework to infer DoA using EEG data from OBA setting.
- We incorporate a meta-learning framework to address cross-subject variability from the OBA.
- We introduce a sequential classification method to reduce the impact of the class-imbalance problem.
- Our proposed Anes-MetaNet achieves the best performance against the other baseline methods.

The remainder of this paper is organized as follows: Section II introduces the proposed method and baseline methods; Section III presents the experimental results and discussion, followed by conclusion in Section IV.

### II. Methodology

In this section, we first introduce the OBA EEG dataset, and the corresponding preprocessing procedure. Then, we discuss the detailed implementation of different models for classification, including CNN, meta-learning CNN (MCNN), LSTM network, as well as our proposed Anes-MetaNet.

#### A. Data description

We use a public office-based anesthesia EEG dataset [25] for this study. The dataset provides EEG recordings of patients during general anesthesia. The induction and maintenance of sedation is done by administering low-dose propofol ramp, intermitted by boluses of propofol, ketamine, dexmedetomidine and lidocaine. Table I lists the provided attributes of the dataset. The RASS score is from a manual assessment of the degree of sedation for patients, ranging from 0 to −5 representing the degree of sedation from consciousness (CON, score 0) to complete loss of consciousness (LOC, score −5). EEG data contains recordings from five channels. In our paper, we test the data of FPZ, FP1 and FP2 channels for the experiment. Due to high motion artifacts during EEG recordings, there are significant noises in the data for a large number of patients.

**TABLE I:** Variable description of the anesthesia data set used in the experiment.

| Variables | Description |
| --- | --- |
| EEG | Raw five channel EEG (bandpass filtered between 0.1-55 Hz). Each row corresponds to channel names in Channelname variable |
| Channelname | EEG channel labels |
| EEG time | Continuous EEG time stamps |
| Fs | Sampling frequency |
| Patient char | Patient characteristics: Procedure type, sex (m/f), age, height (cm), weight (kg) |
| RASS | RASS annotations: 0,-1,-2,-3,-4, and -5 |
| RASS time | Time of RASS assessment |

##### Brain state labels

After visualizing the spectrogram of the EEG data, we find out the SNR is quite low, apparently worse than the HBA data used in [24] and [23]. Hence, it is difficult to classify the OBA data into multiple states as conducted in the previous works. Despite this, the transitioned state between CON and LOC states is critically important for the anesthetists to control the doses of drugs. We then determine to classify the DoA between LOC, CON and the transitioned states in this study. Specifically, we use the EEG epochs with RASS=0 and −1 as the CON state, the EEG epochs with RASS=-2 and −3 as the semi-consciousness (semi-CON) state, and the EEG epochs with RASS=-4 and −5 as LOC state, denoted as *class 0 (CON), class 1 (semi-CON),* and *class 2 (LOC).*

##### Power spectrum analysis

The spectrogram of each subject can be obtained by using multitaper analysis implemented in Chronux Toolbox [37]. The horizontal axis *x_n_* of the spectrogram image represents time in seconds; the vertical axis *y_m_* of the spectrogram represents the frequency in Hz. The value of the ordinate is the signal power in decibel (DB) of *x_i_* time and at *y_j_* frequency. The spectrogram images of 4 subjects are shown in Fig. 1. From the spectrogram images, we visually pick relative high-quality EEG recordings. For example, from Fig. 1, we can find that subjects a and b have good quality EEG data, while the EEG recordings of subjects c and d contain a lot of noises or artifacts.

**Fig. 1:**
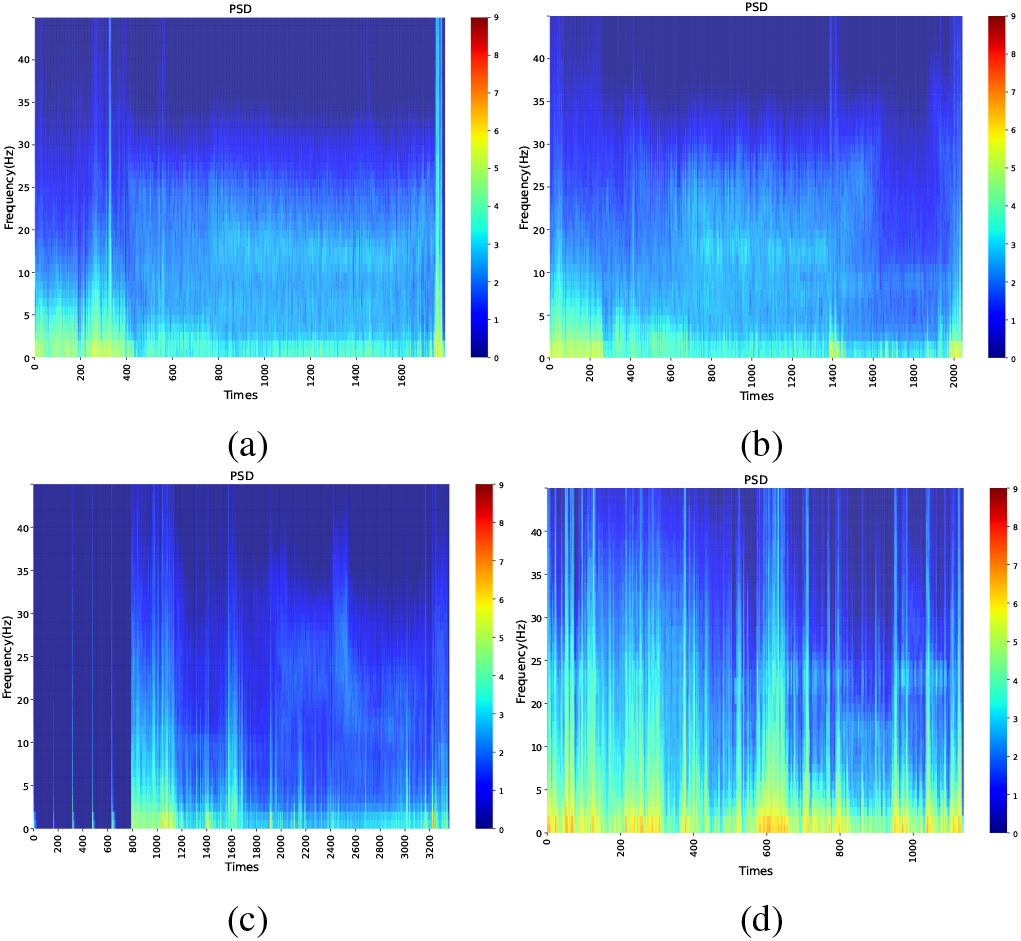
Spectrogram images of 4 subjects: a-d. Compared to subjects a and b, subjects c and d contain significant motion artifacts and noises.

###### 1) Subjects a and b

The middle part of the recording is usually the LOC stage, and the beginning and end parts are usually the CON and semi-CON stages. We name the combination of CON and semi-CON stages as the pre-LOC stage. From subjects a and b, we can observe that the EEG data of LOC state has higher energy in the range of 10-20 Hz. And the EEG data of the pre-LOC state has higher energy within 10 Hz, and lower energy in other frequency ranges. The spectrograms of LOC period and pre-LOC period are significantly different, and satisfactory results can be achieved by classifying LOC and pre-LOC states. However, in the pre-LOC state, there is no obvious difference between the CON and semi-CON states in the spectrogram. As a result, the classification of the CON and semi-CON states is challenging, which motivates us to design a two-stage classification framework.

###### 2) Subject c

From the spectrogram image, we can clearly see that the beginning part of the data of subject c is close to 0. This is caused by technical problems in the recording process of EEG data and we exclude this subject.

###### 3) Subject d

The spectrogram image of subject d shows low SNR of EEG data due to very strong artifacts, and we exclude this subject as well.

By observing the multitaper spectrogram of raw EEG data, we select 13 patients with relatively good data for the following classification experiment.

#### B. Data preprocessing

We use a bandpass filter to filter the EEG data with a frequency band of 0.1 to 40 Hz. The RASS scores are aggregated into classification labels for patients’ anesthesia states. The time intervals between two RASS assessments are not constant, ranging from a few seconds to a few minutes. We add labels for each timestamp of EEG data using the following steps: 1) Find the intermediate time points between all adjacent assessment time periods. 2) If two labels at the beginning time point and ending time point share the same label, then label all the time stamps between two RASS scores as the same label. 3) If the two labels are distinct, label the left half of EEG data within the current interval using the same label as the assessment on the left; label the right half of EEG data with the same label as the assessment on the right. It is worth noting that the labels in 3) are not perfectly accurate. However, we only use them in the training process, and use the labels determined by 2), as the testing data.

#### C. Sequential classification framework

We use a multi-stage sequential classification framework to address the imbalanced data distribution. Taking a three-way classification as an example, in the first step, we observe the data distribution to identify which labeled stage has the largest amount of data and allocate all the rest data to the other class. By using a classification model, such as MCNN, we can obtain a binary classification. In the second step, follow the first-step implementation of classification models on the remaining classes, we can obtain the remaining binary classification result. Given the characteristics of the OBA EEG data, we determine to classify the DoA into three states (CON, semiCON, and LOC) as discussed above. Hence, this classification framework consists of two stages: stage 1: classify pre-LOC vs LOC, and stage 2: classify CON vs semi-CON.

#### D. Convolutional Neural Networks

Given the power spectrum features at one time point is a one dimensional vector, one-dimensional convolutional neural network (1D-CNN) is good to extract the features. As seen in Fig. 2, 1D-CNN includes a convolutional layer, rectified linear unit (ReLU) and a one-dimensional maximum pooling operation. For classification of DoA states, our CNN model is illustrated in Fig. 2, which includes two 1D-CNN blocks and one fully connected block. We define five parameters: let *in_ch_* denote the number of channels for input data, *out_ch_* denote the number of channels for output data, *s_k_* denote the size of the convolution kernels, *p* denote the number of zeros added to each input boundary when performing the convolution operation, and *s_c_* denote the step size of the convolution kernel. Assume that each input feature vector has dimension *n*, the calculation formula of dimension *n*′ of feature vector after a convolution operation is as follows:

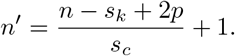

**Fig. 2:**
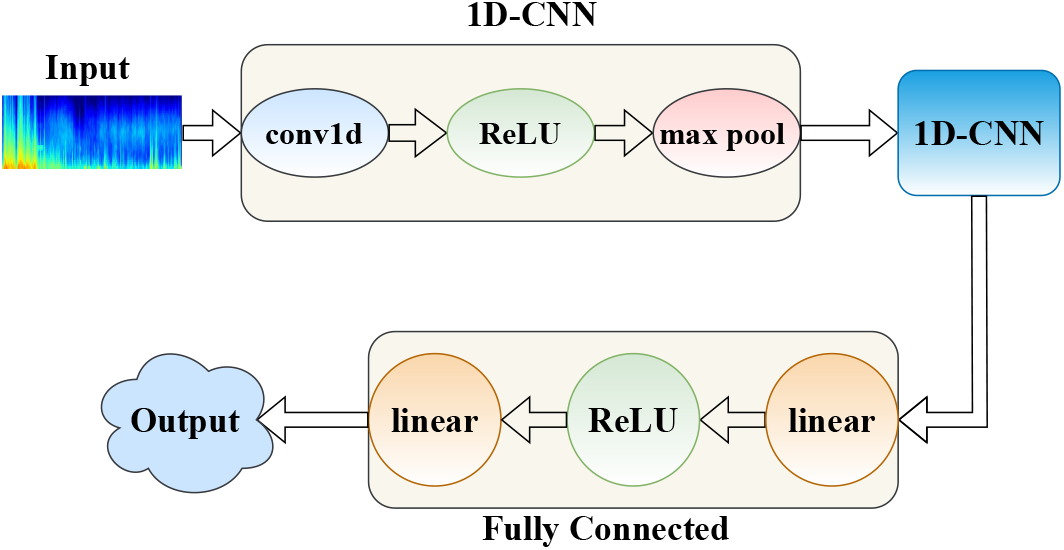
Structure diagram of CNN model.

We define two parameters for the one-dimensional maximum pooling layer. Let *k* denote the window size of the pooling operation and *s_p_* denote the step size of the window of the pooling operation. Assume that the dimension of output feature vectors of one-dimensional convolution is *n*′, the calculation formula of dimension 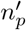 of feature vectors after a maximum pooling operation is as follows:

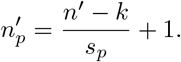

Let *S* denote the total number of input sample. The dimension of the initial input vector is [*S, in_ch_, n*], and the dimension of the output vector after one convolution block operation is 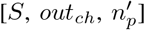.

#### E. Meta-learning CNN

EEG signals vary significantly from subject to subject. The classification function *f* obtained using CNN training from EEG data of some subjects may not be applicable to other subjects. Meta-learning is used in this paper to address this problem. As the meta-learning model is capable of adapting and generalizing to new tasks and new environments that have never been encountered during training time. The difference between meta-learning and supervised learning is that the function of supervised learning directly finds the mapping between features and labels, but the meta-learning function *F*(·) is to find a new *f*, so that the new *f* is suitable for the new machine learning task [35]. The algorithm flow of MCNN is shown as follows:

Step 1: Prepare *N* training tasks. Prepare a test task to evaluate the effects of the parameters learned by MCNN. Prepare a support set and a query set for each task.
Step 2: Define the network structure of CNN and initialize a meta-network parameter *ϕ*^0^. The meta-learning network is the network that is eventually applied to the new test task.
Step 3: Start performing iterative pretraining:

a. For task *i*, assign parameter *ϕ*^0^ of the meta-learning network to the CNN to get 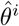.
b. The support set of task *i* is used to optimize and update 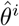 based on the learning rate *α^i^* of task *i*.
c. Calculate loss 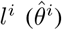 for task *i* using the query set, and compute the gradient of 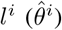 with respect to 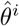.
d. The gradient is multiplied by the learning rate *α^meta^* of the meta-learning network and update *ϕ*^0^ to *ϕ*^1^.
e. Repeat a)-d) on training tasks.
Step 4: Parameters of the meta-learning network are obtained by step 3 and used for testing. Fine-tune the network with the support set of the test task.
Step 5: Evaluate the effectiveness of the MCNN using the query set of the test task.

The structure of the MCNN model is shown in Fig. 3. As noted, the CNN block shown in Fig. 3 is the CNN model presented in Fig. 2. After training, the MCNN model finds a function *F*(·). Whenever a new test task is entered, *F*(·) can always find a set of appropriate parameters to achieve an improved classification performance of CNN model.

**Fig. 3:**
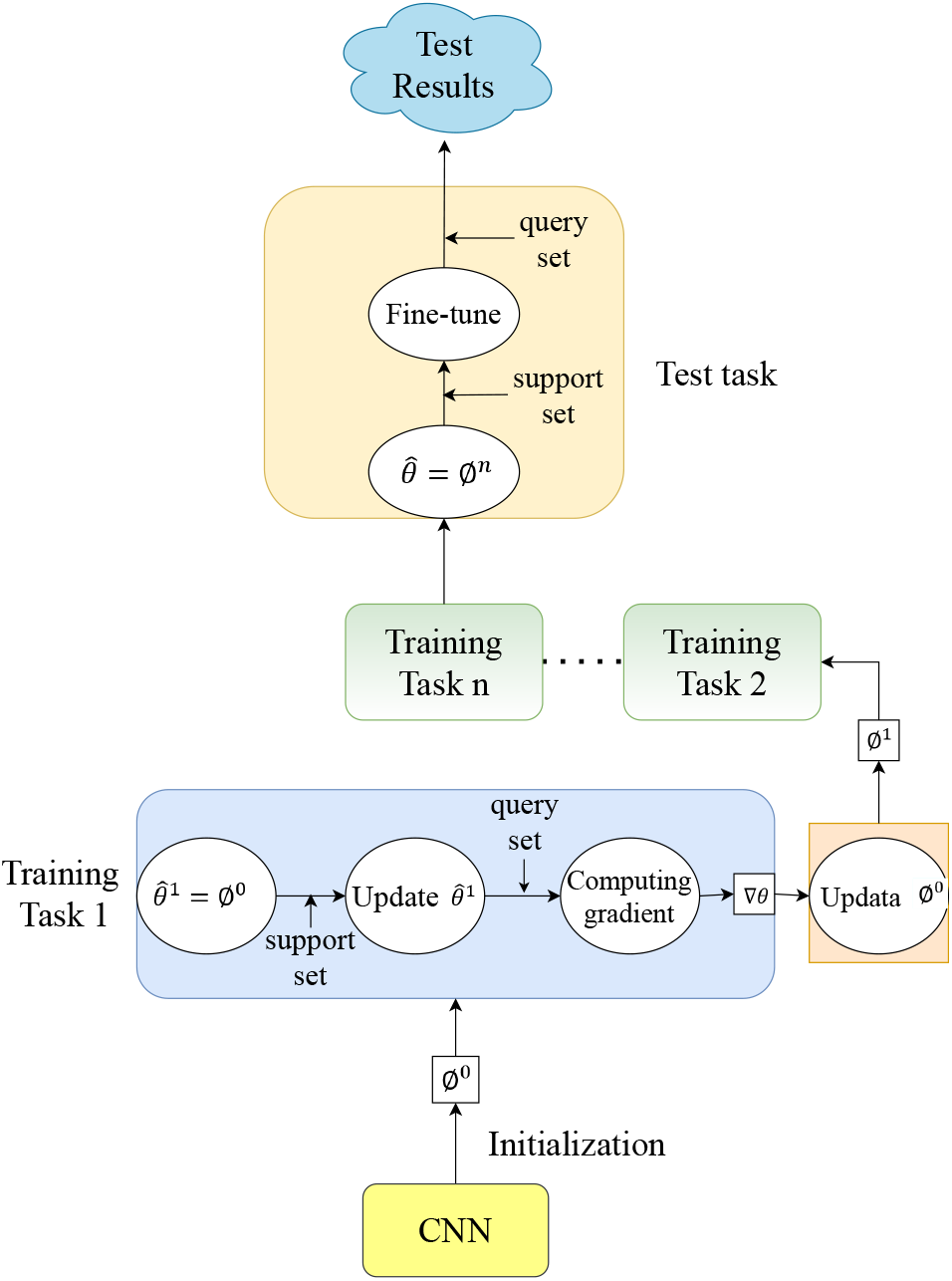
Structure diagram of MCNN model.

#### F. Long Short-Term Memory

Recurrent neural networks (RNN) is good at processing sequential data. LSTM network is a special instance of RNN which mitigates the problems of gradient vanishing and gradient explosion in long sequence training of RNN. LSTM can effectively extract the time characteristics of EEG data, and combine them with the frequency domain features extracted by CNN to achieve high-accuracy classification. LSTM model proposed in this study, consists of two LSTM layers and a fully connected layer. An illustration of the LSTM model is shown in Fig. 4. Let *h*_*t*-1_ denote the output at time *t* – 1, and *x_t_* denote the input at time *t*. Let *c*_*t*–1_ denote the memory at time *t* – 1. *h*_*t*–1_ and *x_t_* are simultaneously input to LSTM layer at time *t*. LSTM has three gates, which are jointly controlled by *h*_*t*–1_ and *x_t_*. The three gates are the forgetting gate, the input gate and the output gate. There are three stages within the LSTM layer:

**Fig. 4:**
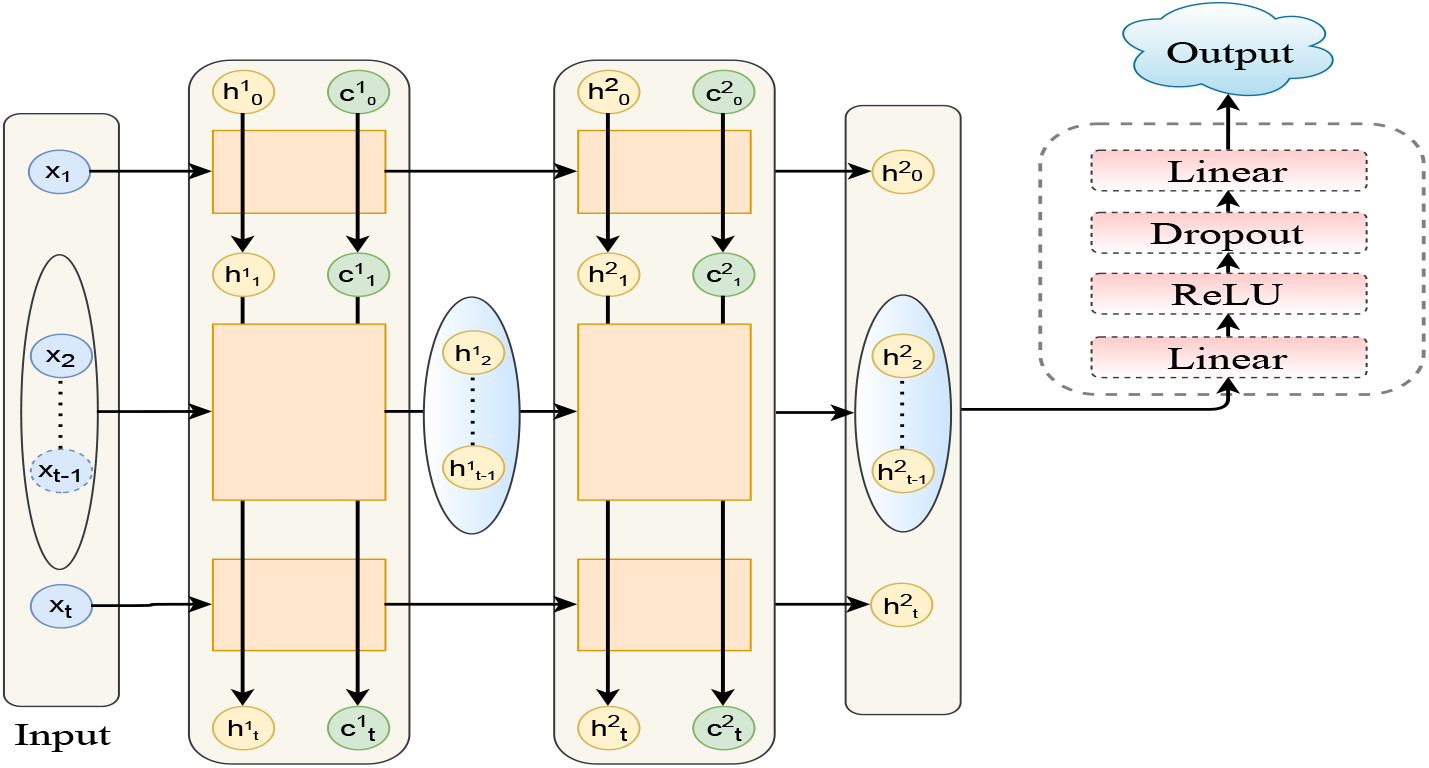
Structure diagram of LSTM model.

##### 1) Forgetting stage

This stage is to selectively forget the input passed in from the previous node. Specifically, the calculated *f_t_* is used as the forgetting gate controller to control the forgetting operation of *c*_*t*–1_ of the previous state. The formula for *f_t_* can be expressed as:

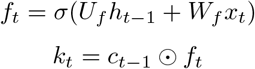

where *U_f_* and *W_f_* are the weight matrix of the forgetting gate, and *σ* is the sigmoid activation function. ⊙ stands for dot multiplication. *k_t_* is the remaining part after forgotten operation of *c*_*t*–1_.

##### 2) Selective memory stage

This stage selectively memorizes the input of this stage. The main purpose is to select and memorize the input *x_t_. i_t_* denotes the input gate controller to control the amount of the input to the memory *k_t_. g_t_* denotes the total input. *c_t_* denotes the newly generated memory. The formula for producing *c_t_* is as follows:

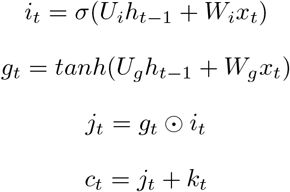

where *U_i_* and *W_i_* are the weight matrix of the input gate, *U_g_* and *W_g_* is the weight matrix used to calculate *g_t_. j_t_* denotes the amount of the input to the memory *k_t_*.

##### 3) Output stage

This stage determines the output of the current state. *o_t_* denotes the output gate controller and *h_t_* denote the output of the current state. The formula for *h_t_* and *o_t_* can be expressed as:

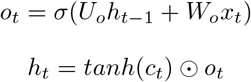

Particularly, in order to prevent over-fitting during neural network training, dropout operations are added to the fully connected block. The parameter of dropout is set to 0.5, which means that neurons at this layer have a random 50% chance of being dropped and not participating in the training during each iteration.

#### G. Proposed Anes-MetaNet

Anes-MetaNet uses CNN and LSTM to capture power spectrum density (PSD) features with the consideration of temporal dependency under the meta-learning training scheme. In Anes-MetaNet, CNN based meta-learning can be trained with multiple stages, depending on the number of classifications. The detailed structure of Anes-MetaNet is shown in Fig. 5. In this study, we classify the EEG anesthetic data into three classes so that two stages are enough. In order to classify CON, semi-CON and LOC, in the first stage of training, labels of EEG data are divided into two categories: Pre-LOC and LOC. We perform the second-stage classification on the correctly classified Pre-LOC data. For Pre-LOC data that is misclassified in the first stage, we eliminate them before the second-stage classification. In the second stage training of Anes-MetaNet, CON and semi-CON states are further classified. We save the features extracted by MCNN as the input of LSTM. The features corresponding to the LOC state are the features extracted from the first trained MCNN model. The features of CON and semi-CON states are the features extracted from the second trained MCNN model. We take the features extracted by CNN for 20 seconds as an input of LSTM model and the prediction of the state in the 20th second as an output. As noted, the meta-learning CNN and LSTM blocks in Fig. 5, are from Fig. 3 and from Fig. 4, respectively.

**Fig. 5:**
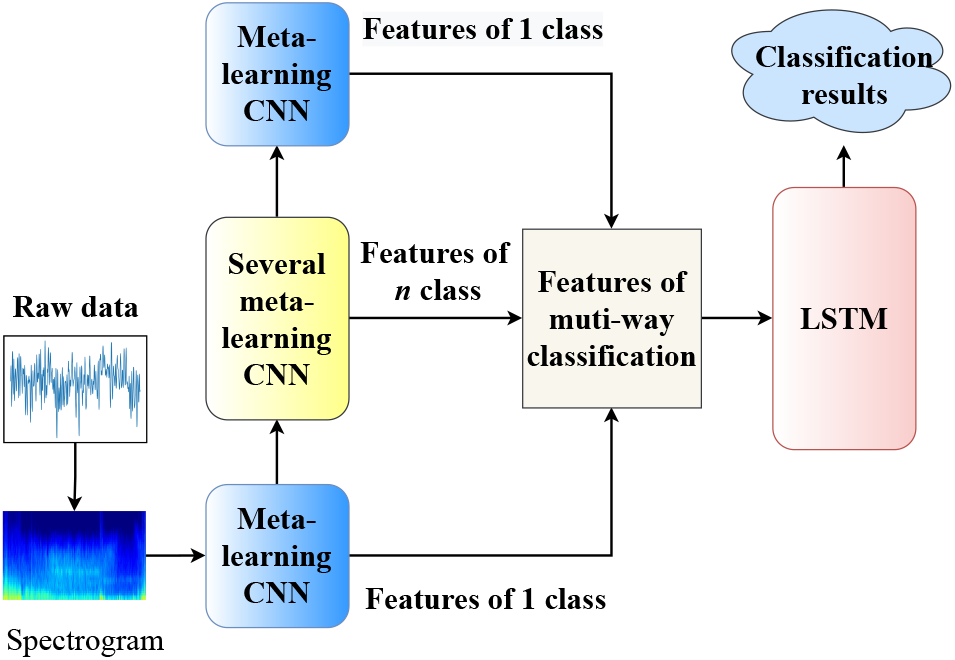
Structure diagram of Anes-MetaNet.

## III. Experiments and Discussion

We conduct seven experiments to demonstrate the effectiveness of our proposed method. Among the 13 subjects, we implement a cross-subject validation by using the data of 10 subjects as the training set, the EEG data of 3 remaining subjects is used as the test data. We take each subject in turn as the training set and the test set for cross validation. In particular, for the MCNN model, we divide the training set and test set with a ratio of 1:1. For each type of label, we randomly extract the same amount of data for training, which can effectively prevent the overfitting problem caused by the problem of class imbalances.

### A. Experiments results

We conduct experiments on three channels of the EEG data. We compare our proposed algorithm with other baseline algorithms, including support vector machines (SVM), random forest (RF), CNN, LSTM, CNN with LSTM (CLSTM), MCNN and Anes-MetaNet. Table II shows the experimental results on classification accuracy and standard deviation of FPZ channel.

**TABLE II:**
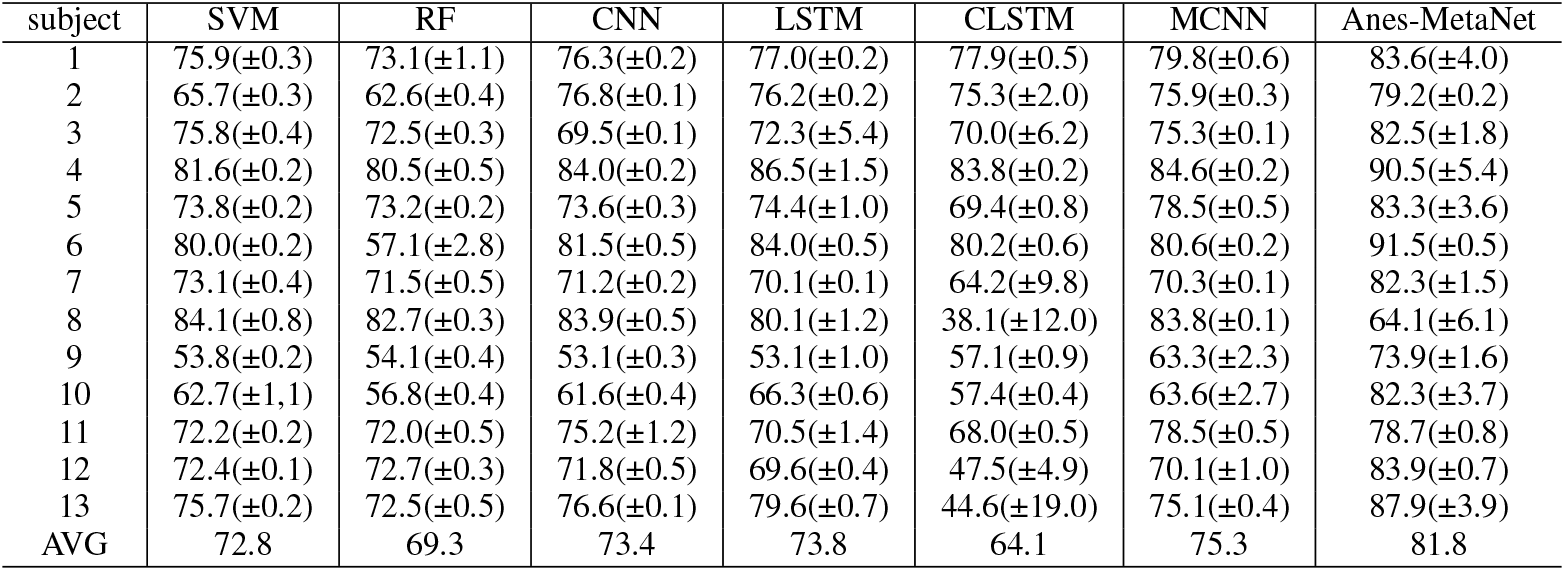
Classification results of EEG data from FPZ channel.

#### Experiments on SVM and RF

Table II shows that the average classification accuracy of SVM and RF is lower than the deep learning method, which demonstrates the superiority of handling large noise under deep learning paradigm. The average classification accuracy of SVM is 72.8%, and the average classification accuracy of RF is 69.3%. SVM uses “gridsearch” algorithm to optimize its hyperparameters, and its classification accuracy is slightly higher than RF’s. The confusion matrices of the classification results of SVM and RF are shown in Fig. 6. As seen in Fig. 6(a), an obvious disadvantage of SVM is its bad performance in classifying the semi-CON state. Fig. 6(b) illustrates that although the RF model can classify the semi-CON state, the accuracy of the classification is less than 30%. Moreover, the classification accuracy of the RF model for the CON state is only about 50%. The above results showcase the difficulty of classifying DoA using traditional machine learning models.

**Fig. 6:**
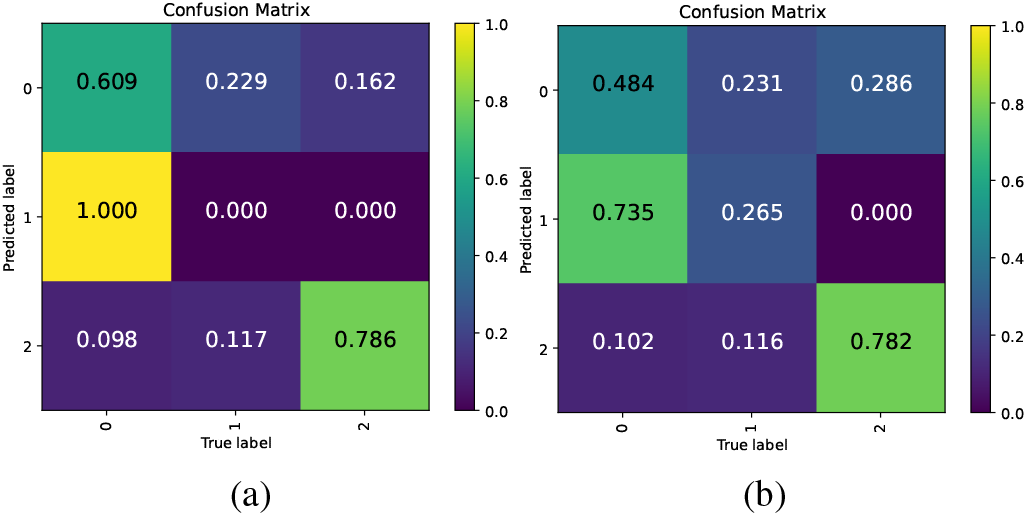
Confusion matrix of the classification results of SVM and RF, where (a) is the confusion matrix of the classification results of SVM, and (b) is the confusion matrix of the classification results of RF. The experimental data is the EEG data of the FP2 channel of subject 1.

#### Experiment on CNN model

In Table II, the average classification accuracy of CNN model has reached 73.4%, which is slightly higher than traditional machine learning methods (SVM and RF), without significant improvement. Fig. 7(a) shows the confusion matrix of the classification results of the CNN model. It can be seen that CNN model also has poor classification performance on semi-CON state. In this experiment, we use the exponential decay method to adjust the learning rate of the CNN model. Fig. 7(b) shows the convergence of loss function of CNN.

**Fig. 7:**
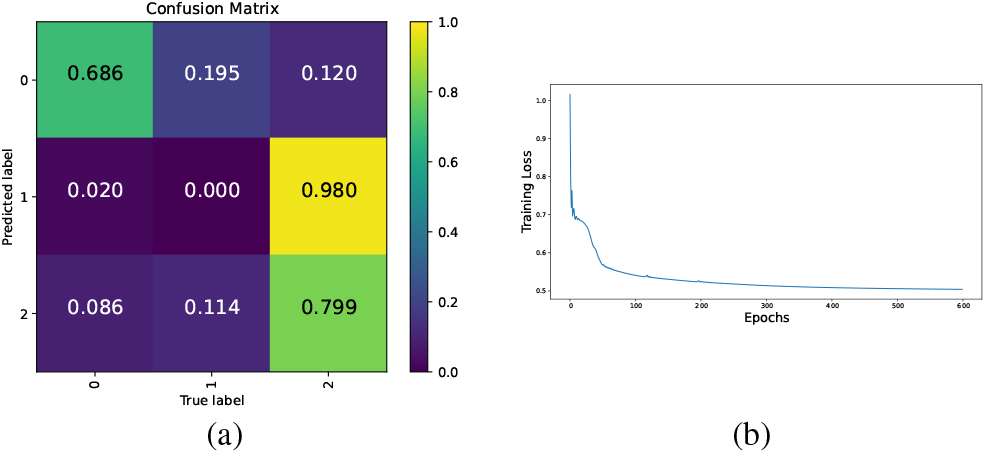
Classification result of CNN model, (a) Confusion matrix diagram of the classification, (b) training loss of CNN.

#### Experiments on LSTM model

In Table II, the results of LSTM model are superior a little to CNN model, with an average classification accuracy of 73.8%. Fig. 8(a) shows that although the accuracy of LSTM model is improved compared with those of SVM and RF models, an obvious shortcoming of LSTM model is that it also cannot classify semi-CON states well. Moreover, Table II illustrates that compared with SVM, RF and CNN models, LSTM model has lower classification stability. Fig. 8(b) shows the loss time series of the LSTM model. It can be seen that the loss curve is gradually stable, which indicates that the model has converged.

**Fig. 8:**
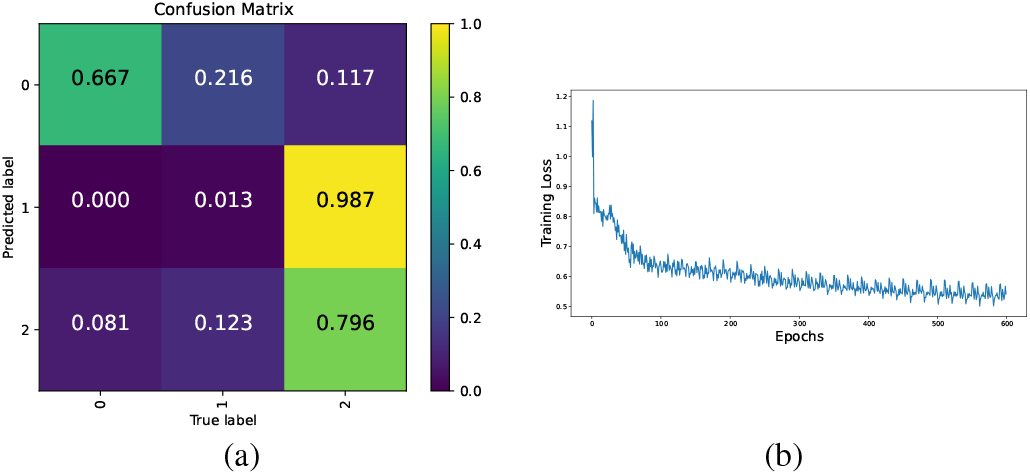
Classification result of LSTM model, (a) Confusion matrix diagram of the classification, (b) training loss of CNN.

**Fig. 9:**
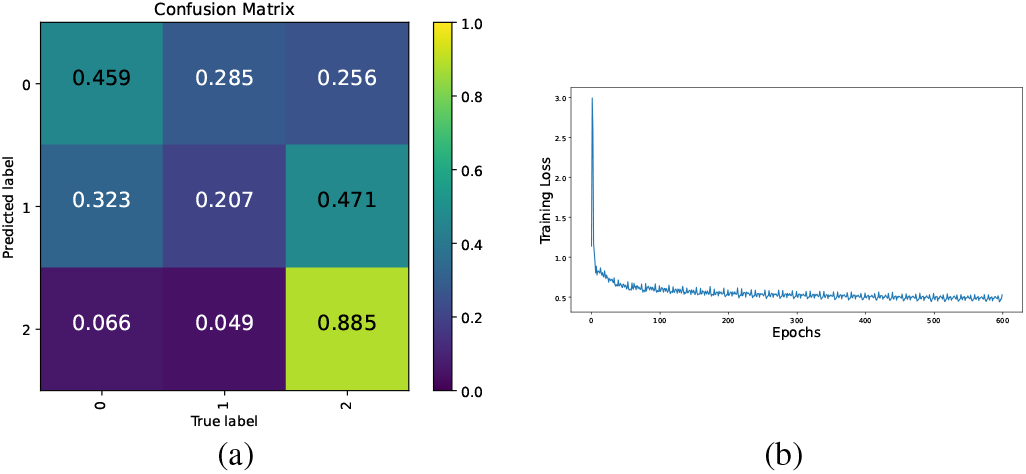
Classification result of CLSTM model, (a) Confusion matrix of the classification by CLSTM, (b) training loss of CLSTM.

#### Experiments on CLSTM model

In this paper, we use CLSTM to represent the model combining CNN and LSTM methods. Table II shows that the classification accuracy of CLSTM is only 64.1%, which is far lower than traditional machine learning models and deep learning models. After analysis, we believe that the poor classification effect of CLSTM is caused by the poor classification results of CNN model. Even though CLSTM model can achieve a higher accuracy on semi-CON state than SVM, CNN and LSTM models; however, the accuracy is only 20.7%. Nevertheless, we still believe CLSTM has a potential to reach a higher classification accuracy once adding some other mechanisms into it. This is confirmed by the following comparative experiments.

#### Experiments on MCNN model

Table II shows that the classification accuracy of the MCNN model has reached 75.3%, which is higher than all traditional machine learning and deep learning models. For poor-quality data such as subject 9, the classification accuracy of MCNN is significantly improved compared to SVM and CNN models. MCNN is trained in two stages, and each stage conducts a binary classification, following the sequential classification method. Fig. 10(a) shows the confusion matrix of the classification results of the first stage. The classification accuracy of the LOC state is about 87.3%, and the classification accuracy of the pre-LOC state is about 78.0%. Fig. 10(b) shows the confusion matrix of the classification results of the second stage. The classification accuracy of the CON state is about 77.8%, and the classification accuracy of the semi-CON state is about 63.2%. Fig. 10(c) shows the confusion matrix of the final classification results of MCNN. The classification accuracy of the CON state is about 58.4%, the classification accuracy of the semi-CON state is about 63.2%, and the classification accuracy of the LOC state is about 87.3%. Fig. 10(c) illustrates that MCNN not only has a high overall classification accuracy, but also can improve the classification accuracy of the CON and Semi-CON states.

**Fig. 10:**
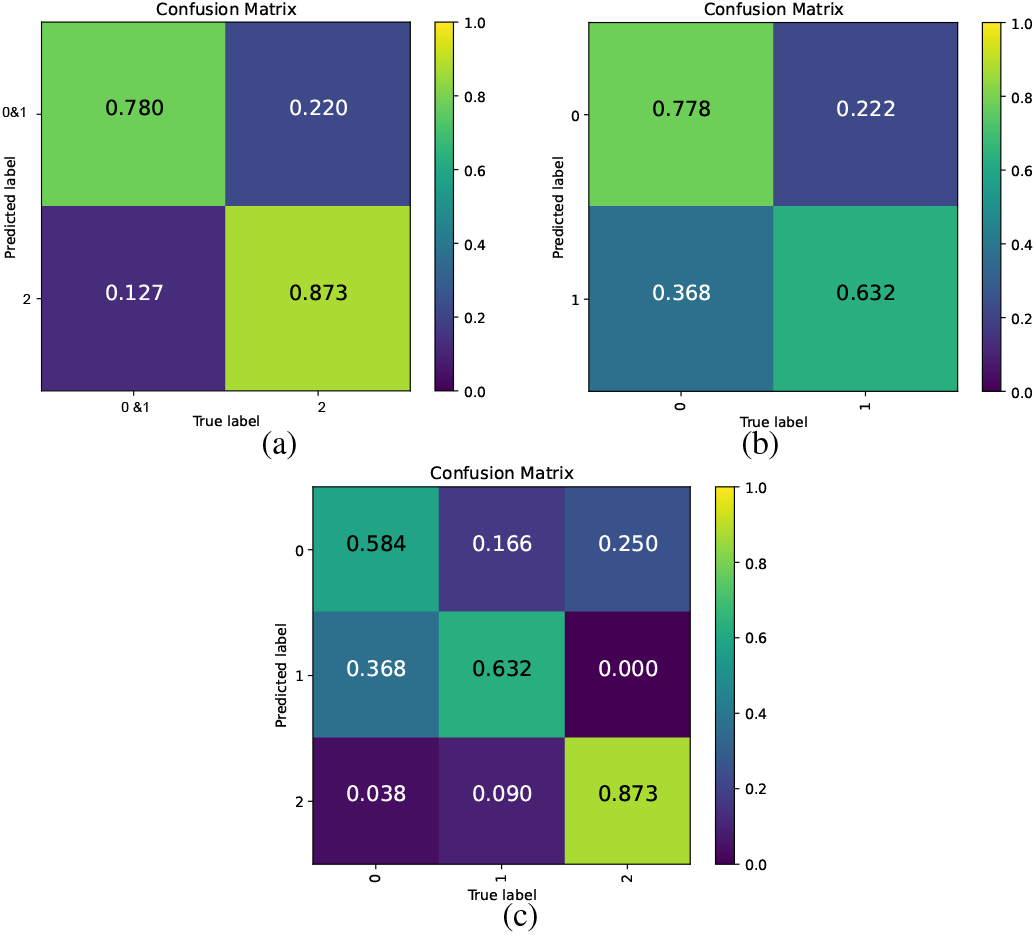
Classification result of MCNN model, (a) confusion matrix of MCNN in the first stage, (b) confusion matrix of MCNN in the second stage, (c) final confusion matrix.

#### Experiments on Anes-MetaNet

Anes-MetaNet achieves an average classification accuracy of 81.8% in the experiments shown in Table II, which is the only model with an average accuracy higher than 80%. For poor quality data such as subjects 9 and 10, the accuracy of Anes-MetaNet is more than 10% higher than other models. Fig. 11(a) shows the confusion matrix of the classification result of Anes-MetaNet. The classification accuracy of CON state is about 64.4%, the classification accuracy of semi-CON state is about 68.9% and LOC state is about 95.1%. Compared to the traditional machine learning models and deep learning models, the proposed Anes-MetaNet can obtain a much higher accuracy on semi-CON state. Furthermore, compared to MCNN model, Anes-MetaNet can have a better classification accuracy on each class. Fig. 11(b) shows the loss curve of Anes-MetaNet during the training process. The curve fluctuates slightly during the training process. This is because we use the dropout layer to prevent the network from overfitting. The loss curve eventually stabilizes, indicating that the network converges. The experimental results of Anes-MetaNet prove that the classification accuracy of the CNN model is an important factor affecting the classification accuracy of the CLSTM model. If the accuracy of the CNN model is low, combining LSTM would not further improve the classification accuracy (see the results using CLSTM). Anes-MetaNet uses meta-learning to improve CNN’s classification ability, which greatly improved LSTM’s classification accuracy.

**Fig. 11:**
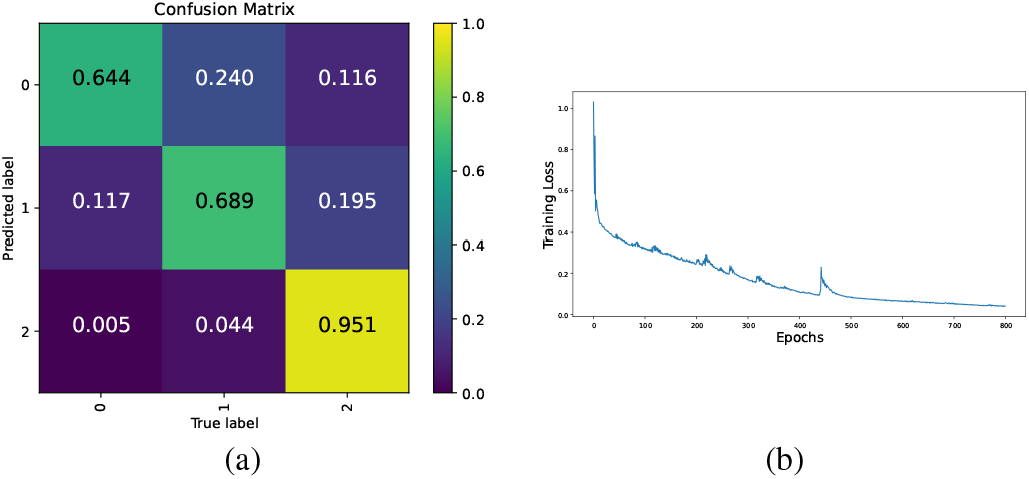
Classification result of Anes-MetaNet model, (a) confusion matrix diagram of the classification, (b) training loss of Anes-MetaNet.

We use t-SNE map to visualize high-level features before the final fully connected layer of CNN, and the result is illustrated in Fig. 12. As seen, class 2 (LOC) has a more pronounced difference from other classes (0: CON and 1: semi-CON), which justifies our previous choice of using a two-stage training scheme to first distinguish LOC from pre-LOC (combining CON and semi-CON). Besides this, we can see that class 0 and class 1 are mixed together closely, which explains why the traditional machine learning models and deep learning models cannot classify the semi-CON state well.

**Fig. 12:**
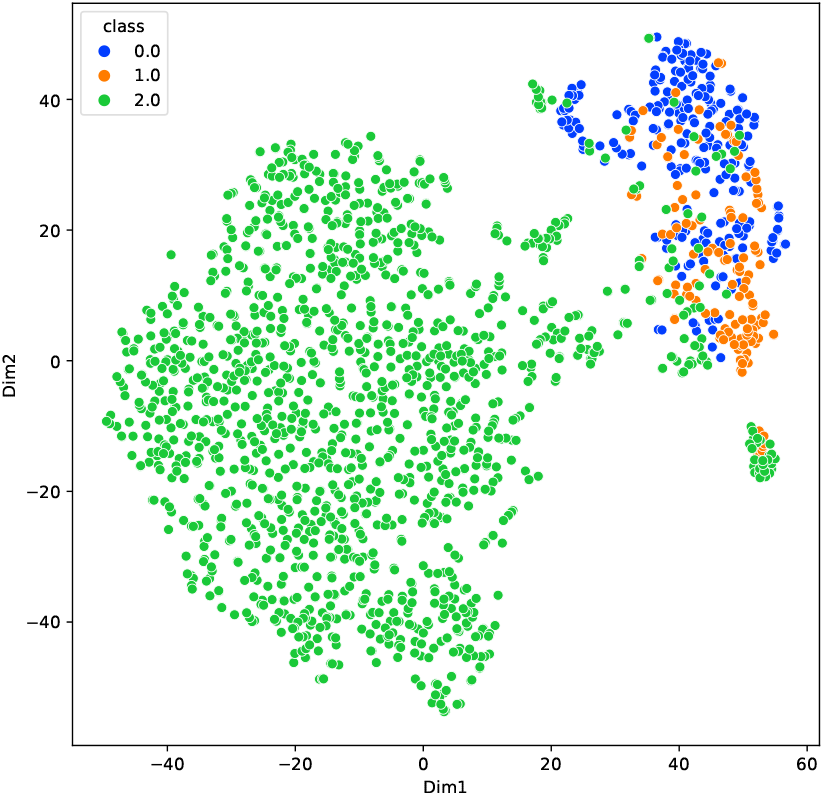
Visualization of t-SNE maps.

Table III and Table IV respectively show the experimental results of the EEG data recorded by the FP1 and FP2 channels of 13 subjects. By comparing the results in Tables II-IV, we can draw the following conclusions:

1. As the OBA EEG data contains a high level of noises, traditional machine learning and deep learning models usually cannot perform well; however, meta-learning is a good method to reduce the cross-subject noises and improve the classification accuracy by combining with deep learning methods.
2. When using single-channel EEG data, the data of FP1 and FP2 channels are better for the classification usage of brain states than the data of FPZ channel (see the average results of all methods in Tables II-IV). As noted, Anes-MetaNet achieved the highest accuracy for classifying the DoA using FP1 channel of the EEG data.
3. LSTM is easy to cause the instability for the classification results (see the results using LSTM, CLSTM and Anes-MetaNet). Nevertheless, LSTM improve the classification accuracy by comparing the results between using MCNN and Anes-MetaNet.

**TABLE III:** Experimental results of EEG data from FP1 channel.

| Subject | SVM | RF | CNN | LSTM | CLSTM | MCNN | Anes-MetaNet |
| --- | --- | --- | --- | --- | --- | --- | --- |
| 1 | 80.0(±0.1) | 80.5(±0.1) | 81.2(±0.2) | 87.5(±0.5) | 86.4(±1.3) | 86.2(±0.9) | 86.4(±1.0) |
| 2 | 75.5(±0.1) | 69.9(±0.1) | 76.3(±0.1) | 66.2(±2.9) | 70.5(±2.7) | 77.6(±0.2) | 81.3(±0.2) |
| 3 | 71.6(±0.2) | 69.4(±0.1) | 68.9(±1.5) | 79.8(±0.1) | 70.4(±2.2) | 78.5(±0.9) | 78.5(±2.8) |
| 4 | 88.9(±0.1) | 91.1(±0.1) | 86.4(±2.7) | 90.5(±0.5) | 87.9(±1.4) | 89.8(±0.8) | 92.8(±3.3) |
| 5 | 72.0(±0.4) | 72.4(±0.1) | 73.6(±1.9) | 75.3(±0.1) | 72.4(±0.4) | 78.7(±0.2) | 79.3(±1.9) |
| 6 | 82.2(±0.2) | 80.6(±0.1) | 81.8(±0.1) | 84.9(±0.3) | 83.4(±2.1) | 84.0(±0.3) | 90.5(±0.1) |
| 7 | 73.0(±0.1) | 72.5(±0.1) | 74.0(±0.1) | 76.5(±0.1) | 76.6(±0.2) | 74.8(±0.3) | 89.5(±0.8) |
| 8 | 85.4(±0.6) | 83.5(±0.1) | 84.0(±1.2) | 82.9(±2.4) | 55.1(±15.9) | 84.0(±0.4) | 86.8(±0.9) |
| 9 | 52.1(±0.3) | 54.7(±0.3) | 54.0(±0.7) | 64.5(±1.3) | 62.7(±2.1) | 63.6(±0.6) | 71.9(±0.8) |
| 10 | 58.5(±0.3) | 59.0(±0.5) | 58.8(±0.8) | 49.7(±2.7) | 54.3(±0.2) | 57.1(±0.4) | 74.1(±0.8) |
| 11 | 76.3(±0.2) | 77.1(±0.1) | 75.3(±0.2) | 64.4(±0.2) | 66.2(±1.7) | 81.7(±1.1) | 84.7(±2.1) |
| 12 | 76.4(±0.2) | 76.1(±0.1) | 75.3(±1.0) | 79.1(±1.3) | 75.5(±0.5) | 74.7(±0.3) | 83.9(±0.5) |
| 13 | 76.2(±0.1) | 76.2(±0.2) | 75.6(±0.1) | 75.6(±1.7) | 77.2(±0.4) | 75.8(±0.3) | 81.6(±2.9) |
| AVG | 74.4 | 74.0 | 74.2 | 75.1 | 72.2 | 77.4 | 83.1 |

**TABLE IV:**
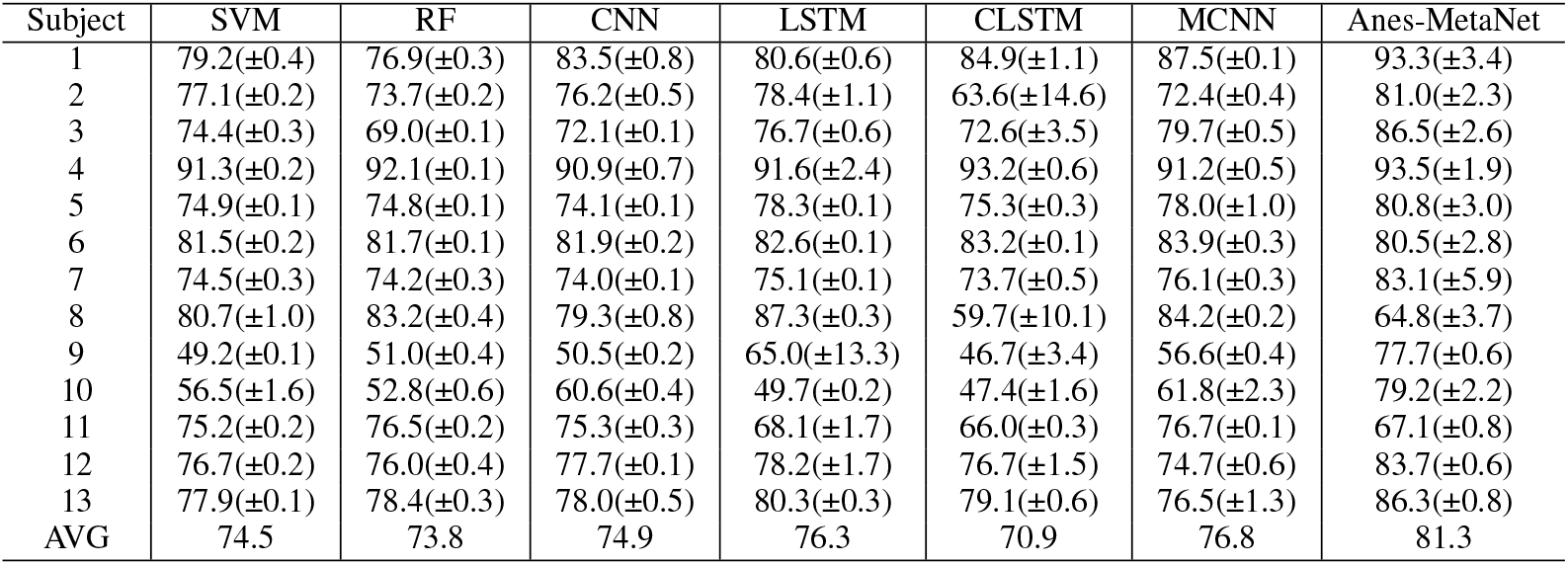
Experimental results base on EEG data from FP2 channel.

| Subject | SVM | RF | CNN | LSTM | CLSTM | MCNN | Anes-MetaNet |
| --- | --- | --- | --- | --- | --- | --- | --- |
| 1 | 79.2(±0.4) | 76.9(±0.3) | 83.5(±0.8) | 80.6(±0.6) | 84.9(±1.1) | 87.5(±0.1) | 93.3(±3.4) |
| 2 | 77.1(±0.2) | 73.7(±0.2) | 76.2(±0.5) | 78.4(±1.1) | 63.6(±14.6) | 72.4(±0.4) | 81.0(±2.3) |
| 3 | 74.4(±0.3) | 69.0(±0.1) | 72.1(±0.1) | 76.7(±0.6) | 72.6(±3.5) | 79.7(±0.5) | 86.5(±2.6) |
| 4 | 91.3(±0.2) | 92.1(±0.1) | 90.9(±0.7) | 91.6(±2.4) | 93.2(±0.6) | 91.2(±0.5) | 93.5(±1.9) |
| 5 | 74.9(±0.1) | 74.8(±0.1) | 74.1(±0.1) | 78.3(±0.1) | 75.3(±0.3) | 78.0(±1.0) | 80.8(±3.0) |
| 6 | 81.5(±0.2) | 81.7(±0.1) | 81.9(±0.2) | 82.6(±0.1) | 83.2(±0.1) | 83.9(±0.3) | 80.5(±2.8) |
| 7 | 74.5(±0.3) | 74.2(±0.3) | 74.0(±0.1) | 75.1(±0.1) | 73.7(±0.5) | 76.1(±0.3) | 83.1(±5.9) |
| 8 | 80.7(±1.0) | 83.2(±0.4) | 79.3(±0.8) | 87.3(±0.3) | 59.7(±10.1) | 84.2(±0.2) | 64.8(±3.7) |
| 9 | 49.2(±0.1) | 51.0(±0.4) | 50.5(±0.2) | 65.0(±13.3) | 46.7(±3.4) | 56.6(±0.4) | 77.7(±0.6) |
| 10 | 56.5(±1.6) | 52.8(±0.6) | 60.6(±0.4) | 49.7(±0.2) | 47.4(±1.6) | 61.8(±2.3) | 79.2(±2.2) |
| 11 | 75.2(±0.2) | 76.5(±0.2) | 75.3(±0.3) | 68.1(±1.7) | 66.0(±0.3) | 76.7(±0.1) | 67.1(±0.8) |
| 12 | 76.7(±0.2) | 76.0(±0.4) | 77.7(±0.1) | 78.2(±1.7) | 76.7(±1.5) | 74.7(±0.6) | 83.7(±0.6) |
| 13 | 77.9(±0.1) | 78.4(±0.3) | 78.0(±0.5) | 80.3(±0.3) | 79.1(±0.6) | 76.5(±1.3) | 86.3(±0.8) |
| AVG | 74.5 | 73.8 | 74.9 | 76.3 | 70.9 | 76.8 | 81.3 |

From the above experimental results, it can be seen that the classification results of CNN and LSTM are better than SVM and RF. However, the result of CLSTM is not as good as CNN or LSTM. The reason is that the classification results using CNN are not good enough (no better or just slightly better than a random guess); thus, adding LSTM makes the prediction further away from the true label given too many wrongly classified samples by CNN. The training of LSTM needs better represented features and predicted labels rendered by CNN. The wrong features and labels lead to the inability of LSTM to train an accurate model. The classification results of MCNN are not only better than CNN in the final accuracy, but also much better than CNN in the classification of the more difficult task (CON vs semi-CON). From the confusion matrix, we can see that MCNN achieves a high accuracy in the classification results of each class, while CNN only has high accuracy in the classification of LOC state and CON state, and the accuracy in the classification of semi-CON state is about 0. Such a phenomenon can be explained that metalearning and two-stage classification mechanism in MCNN are much helpful to improve the classification capability of CNN, so that the features extracted by the CNN model can more accurately represent the DoA. Compared CLSTM to Anes-MetaNet, accurately representing the input features for LSTM is important for the further feature learning in improving the classification accuracy.

## IV. Conclusion

In this study, we propose an Anes-MetaNet to solve the DoA classification problem in office-based environments, in which, studies on the classification of anesthesia EEG data have not been well investigated. The model we develop, provides good performance for the classification of brain states using anesthesia EEG data. The proposed Anes-MetaNet can mitigate the individual differences among subjects. We train the model in a sequential way to reach a better classification accuracy. The t-SNE visualization validates our two-stage training mechanism. In the future, a systematic comparison of OBA EEG data and HBA EEG data under different anesthetics need to be conducted with regards to model accuracy and interpretability.

## V. Acknowledge

Q. Wang and Y. Chen are partly supported by National Natural Science Foundation of China (Grant No. 72101066, 72131005, 91846301).

## References

[1] E.N. Brown, P. L. Purdon, and C. J. Van Dort, “General anesthesia and altered states of arousal: a systems neuroscience analysis,” Annual review of neuroscience, vol. 34, pp. 601–628, 2011.

[2] B. A. Fritz, P. L. Kalarickal, H. R. Maybrier, M. R. Muench, D. Dearth, Y. Chen, K. E. Escallier, A. B. Abdallah, N. Lin, and M. S. Avidan, “Intraoperative electroencephalogram suppression predicts postoperative delirium,” Anesthesia and analgesia, vol. 122, no. 1, p. 234, 2016.

[3] P. L. Purdon, E. T. Pierce, E. A. Mukamel, M. J. Prerau, J. L. Walsh, K. F. K. Wong, A. F. Salazar-Gomez, P. G. Harrell, A. L. Sampson, A. Cimenser, et al., “Electroencephalogram signatures of loss and recovery of consciousness from propofol,” Proceedings of the National Academy of Sciences, vol. 110, no. 12, pp. E1142–E1151, 2013.

[4] S. Chakravarty, J. A. Donoghue, A. S. Waite, M. Mahnke, I. C. Garwood, E. K. Miller, and E. N. Brown, “Closed-loop control of anesthetic state in non-human primates,” bioRxiv, 2021.

[5] C. N. Sessler, M. S. Gosnell, M. J. Grap, G. M. Brophy, P. V. O’Neal, K. A. Keane, E. P. Tesoro, and R. Elswick, “The richmond agitation– sedation scale: validity and reliability in adult intensive care unit patients,” American journal of respiratory and critical care medicine, vol. 166, no. 10, pp. 1338–1344, 2002.

[6] T. A. Bowdle, “Depth of anesthesia monitoring,” Anesthesiology Clinics of North America, vol. 24, no. 4, pp. 793–822, 2006.

[7] X.-S. Zhang, R. J. Roy, and E. W. Jensen, “EEG complexity as a measure of depth of anesthesia for patients,” IEEE transactions on biomedical engineering, vol. 48, no. 12, pp. 1424–1433, 2001.

[8] F. L. da Silva, “EEG and MEG: relevance to neuroscience,” Neuron, vol. 80, no. 5, pp. 1112–1128, 2013.

[9] O. Akeju, A. H. Song, A. E. Hamilos, K. J. Pavone, F. J. Flores, E. N. Brown, and P. L. Purdon, “Electroencephalogram signatures of ketamine anesthesia-induced unconsciousness,” Clinical neurophysiology, vol. 127, no. 6, pp. 2414–2422, 2016.

[10] R. Shalbaf, H. Behnam, and H. J. Moghadam, “Monitoring depth of anesthesia using combination of EEG measure and hemodynamic variables,” Cognitive Neurodynamics, vol. 9, no. 1, pp. 41–51, 2015.

[11] M. Jospin, P. Caminal, E. W. Jensen, H. Litvan, M. Vallverdü, M. M. Struys, H. E. Vereecke, and D. T. Kaplan, “Detrended fluctuation analysis of EEG as a measure of depth of anesthesia,” IEEE transactions on biomedical engineering, vol. 54, no. 5, pp. 840–846, 2007.

[12] T. Nguyen-Ky, P. Wen, and Y. Li, “Consciousness and depth of anesthesia assessment based on bayesian analysis of EEG signals,” IEEE Transactions on Biomedical Engineering, vol. 60, no. 6, pp. 1488–1498, 2013.

[13] R. Shalbaf, H. Behnam, J. W. Sleigh, A. Steyn-Ross, and L. J. Voss, “Monitoring the depth of anesthesia using entropy features and an artificial neural network,” Journal of neuroscience methods, vol. 218, no. 1, pp. 17–24, 2013.

[14] M. Peker, A. Arslan, B. Sen, F. V. Çelebi, and A. But, “A novel hybrid method for determining the depth of anesthesia level: Combining relieff feature selection and random forest algorithm (relieff+ rf),” in 2015 International Symposium on Innovations in Intelligent SysTems and Applications (INISTA), pp. 1–8, IEEE, 2015.

[15] M. Jahanseir, K. Setarehdan, and S. Momenzadeh, “Estimation of the depth of anesthesia by using a multioutput least-square support vector regression,” Turkish Journal of Electrical Engineering & Computer Sciences, vol. 26, no. 6, pp. 2792–2801, 2018.

[16] W. Saadeh, F. H. Khan, and M. A. B. Altaf, “Design and implementation of a machine learning based EEG processor for accurate estimation of depth of anesthesia,” IEEE transactions on biomedical circuits and systems, vol. 13, no. 4, pp. 658–669, 2019.

[17] K. He, X. Zhang, S. Ren, and J. Sun, “Identity mappings in deep residual networks,” in European conference on computer vision, pp. 630–645, Springer, 2016.

[18] S. Zhou, X. Li, Y. Chen, S. T. Chandrasekaran, and A. Sanyal, “Temporal-coded deep spiking neural network with easy training and robust performance,” arXiv preprint arXiv:1909.10837, 2019.

[19] T. Young, D. Hazarika, S. Poria, and E. Cambria, “Recent trends in deep learning based natural language processing,” IEEE Computational intelligence Magazine, vol. 13, no. 3, pp. 55–75, 2018.

[20] Y. Fujisawa, Y. Otomo, Y. Ogata, Y. Nakamura, R. Fujita, Y. Ishitsuka, R. Watanabe, N. Okiyama, K. Ohara, and M. Fujimoto, “Deep-learningbased, computer-aided classifier developed with a small dataset of clinical images surpasses board-certified dermatologists in skin tumour diagnosis,” British Journal of Dermatology, vol. 180, no. 2, pp. 373–381, 2019.

[21] M. Jiao, D. Wang, Y. Yang, and F. Liu, “More intelligent and robust estimation of battery state-of-charge with an improved regularized extreme learning machine,” Engineering Applications of Artificial Intelligence, vol. 104, p. 104407, 2021.

[22] Y. Park, S.-H. Han, W. Byun, J.-H. Kim, H.-C. Lee, and S.-J. Kim, “A real-time depth of anesthesia monitoring system based on deep neural network with large edo tolerant EEG analog front-end,” IEEE Transactions on Biomedical Circuits and Systems, vol. 14, no. 4, pp. 825–837, 2020.

[23] R. Li, Q. Wu, J. Liu, Q. Wu, C. Li, and Q. Zhao, “Monitoring depth of anesthesia based on hybrid features and recurrent neural network,” Frontiers in Neuroscience, vol. 14, p. 26, 2020.

[24] S. Afshar, R. Boostani, and S. Saeid, “A combinatorial deep learning structure for precise depth of anesthesia estimation from EEG signals,” IEEE Journal of Biomedical and Health Informatics, 2021.

[25] S. B. Nagaraj, P. L. Purdon, F. Shapiro, B. Westover, et al., “Electroencephalogram monitoring of depth of anesthesia during office-based anesthesia,” bioRxiv, 2020.

[26] T. Zoughi and R. Boostani, “Presenting a combinatorial feature to estimate depth of anesthesia,” International Journal of Signal Processing, vol. 6, no. 2, pp. 10–14, 2010.

[27] S. Afrasiabi, R. Boostani, S. Koochaki, and F. Zand, “Presenting an effective EEG-based index to monitor the depth of anesthesia,” in The 16th CSI International Symposium on Artificial Intelligence and Signal Processing (AISP 2012), pp. 557–562, IEEE, 2012.

[28] K. Polat and S. Güneş, “Classification of epileptiform EEG using a hybrid system based on decision tree classifier and fast fourier transform,” Applied Mathematics and Computation, vol. 187, no. 2, pp. 1017–1026, 2007.

[29] H. Cecotti and A. Graeser, “Convolutional neural network with embedded fourier transform for EEG classification,” in 2008 19th International Conference on Pattern Recognition, pp. 1–4, IEEE, 2008.

[30] B. Babadi and E. N. Brown, “A review of multitaper spectral analysis,” IEEE Transactions on Biomedical Engineering, vol. 61, no. 5, pp. 1555–1564, 2014.

[31] B. Noureddin, P. D. Lawrence, and G. E. Birch, “Online removal of eye movement and blink eeg artifacts using a high-speed eye tracker,” IEEE Transactions on Biomedical Engineering, vol. 59, no. 8, pp. 2103–2110, 2011.

[32] U. R. Acharya, S. L. Oh, Y. Hagiwara, J. H. Tan, and H. Adeli, “Deep convolutional neural network for the automated detection and diagnosis of seizure using EEG signals,” Computers in biology and medicine, vol. 100, pp. 270–278, 2018.

[33] V. J. Lawhern, A. J. Solon, N. R. Waytowich, S. M. Gordon, C. P. Hung, and B. J. Lance, “Eegnet: a compact convolutional neural network for EEG-based brain–computer interfaces,” Journal of Neural Engineering, vol. 15, no. 5, p. 056013, 2018.

[34] Z. Jiao, X. Gao, Y. Wang, J. Li, and H. Xu, “Deep convolutional neural networks for mental load classification based on EEG data,” Pattern Recognition, vol. 76, pp. 582–595, 2018.

[35] R. Vilalta and Y. Drissi, “A perspective view and survey of metalearning,” Artificial intelligence review, vol. 18, no. 2, pp. 77–95, 2002.

[36] J. Vanschoren, “Meta-learning: A survey,” arXiv preprint arXiv:1810.03548, 2018.

[37] H. Bokil, P. Andrews, J. E. Kulkarni, S. Mehta, and P. P. Mitra, “Chronux: a platform for analyzing neural signals,” Journal of neuroscience methods, vol. 192, no. 1, pp. 146–151, 2010.

